# MetaFX: feature extraction from whole-genome metagenomic sequencing data

**DOI:** 10.1101/2025.05.27.656413

**Authors:** Artem Ivanov, Vladimir Popov, Maxim Morozov, Evgenii Olekhnovich, Vladimir Ulyantsev

## Abstract

**Motivation:** Microbial communities consist of thousands of microorganisms and viruses and have a tight connection with an environment, such as gut microbiota modulation of host body metabolism. However, the direct relationship between the presence of certain microorganism and host state often remains unknown. Toolkits using reference-based approaches are limited to microbes present in databases. Reference-free methods often require enormous resources for metagenomic assembly or results in many poorly interpretable features based on k-mers.

**Results:** Here we present MetaFX – an open-source library for feature extraction from whole-genome metagenomic sequencing data and classification of groups of samples. Using a large volume of metagenomic samples deposited in databases, MetaFX compares samples grouped by metadata criteria (e.g. disease, treatment, etc) and constructs genomic features distinct for certain types of communities. Features constructed based on statistical k-mer analysis and de Bruijn graphs partition. Those features are used in machine learning models for classification of novel samples. Extracted features can be visualised on de Bruijn graphs and annotated for providing biological insights. We demonstrate the utility of MetaFX by building classification models for 590 human gut samples with inflammatory bowel disease. Our results outperform the previous research disease prediction accuracy up to 17%, and improves classification results compared to taxonomic analysis by 9 *±* 10% on average.

**Availability:** MetaFX is a feature extraction toolkit applicable for metagenomic datasets analysis and samples classification. The source code, test data, and relevant information for MetaFX are freely accessible at https://github.com/ctlab/metafx under the MIT License.

## Introduction

Microbial communities inhabit diverse ecological niches, including soil, water reservoirs, the human gut and others (Garner et al. [2023], Fierer [2017], Human Microbiome Project Consortium [2012]). Metagenomic sequencing data analysis is a well-established approach to analyse the structure and functions of microbial communities. For example, many ongoing studies are exploring the gut microbiome with regards to health and disease, as well as contributing to the development of diagnostic and therapeutic strategies (Olekhnovich et al. [2023], Ivanova et al. [2022], Olekhnovich et al. [2021], Lloyd-Price et al. [2019], Jie et al. [2017], Yu et al. [2017], Qin et al. [2012]). However, existing metagenomic analysis methods, including taxonomic and functional annotation or genome-resolved metagenomics, provide only limited resolution of microbiome properties.

Here, we present MetaFX, an open-source library for the extraction of features from metagenomic sequencing data and the classification of samples’ groups. Its main aim is to construct features initially in a reference-free manner, while keeping the possibility for their consequent analysis and annotation. The tool takes as an input metagenomic data in fastq or fasta format from short-read Illumina sequencing platforms and constructs features in the form of contigs or branching paths in the de Bruijn graph. These features can be further analysed for biological significance (e.g. taxonomic or functionally annotated) as well as be used to train machine learning models for classification of novel samples.

## Software and implementation

### Concept of MetaFX

MetaFX is a command line tool optimised for multi-threaded environments, which makes it possible to process large metagenomic datasets (hundreds of samples) on computational servers within a reasonable time limit (for example, 220 samples were successfully processed in 40 hours with 32 threads and 400Gb RAM). The outline of MetaFX library is presented in Figure 1. MetaFX is implemented as a pipeline, which calls several tools for feature extraction and visualisations, and custom python scripts for data analysis and manipulation. Most of the feature extraction modules are based on the MetaFast (Ulyantsev et al. [2016]) toolkit, which was substantially modified by creating new classes for supervised feature extraction. The idea for MetaFX raised from international challenge on inflammatory bowel disease prediction based on patients gut microbiome, that authors of this paper won (Khachatryan et al. [2023]).

**Fig. 1.**
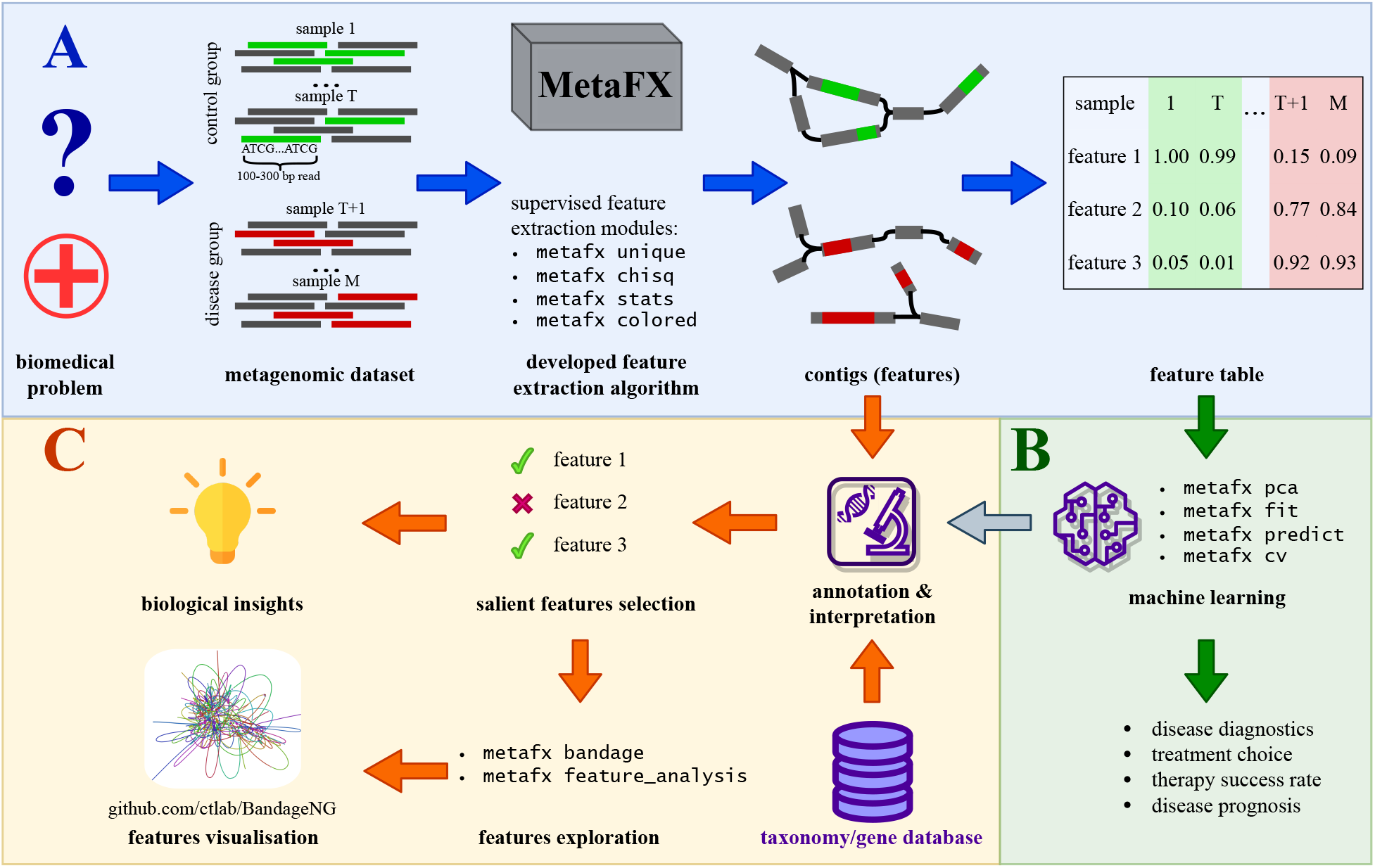
Overall pipeline of data analysis from raw reads to interpretable features and classification model in MetaFX. (A) Feature extraction steps; (B) classification model training steps; (C) features annotation and visualisation steps.

Conceptually, all MetaFX modules can be divided into three categories (Supplementary Figure 1). Two of them aimed at constructing distinct features from samples. Features can be extracted jointly from all of the input metagenomes in case of the lack of associated samples information – these methods are described in unsupervised methods section. Alternatively, all samples can be split into several groups based on samples’ metadata information. In that case, supervised methods are applied to extract features for each group independently. The last group of methods encompass various scripts for subsequent analysis of extracted features. It includes classification model training and prediction, preparation for features visualisation, and also utilities for computations speed up and formats transformations.

### Unsupervised methods for feature extraction

Unsupervised methods provides the ability to extract features for all input samples. It can be useful for exploratory data analysis and samples clustering when there is no *a priori* division of metagenomes into categories. Two such methods have been implemented.

MetaFX `metafast` module utilises original MetaFast algorithm of constructing condensed de Bruijn graph of all samples and then its iterative breakage into connected components based on coverage thresholds (see details in Ulyantsev et al. [2016]). Each component then used as a single feature. Users can tune the features by the input parameters controlling the size of components and coverage thresholds.

Another module `metaspades` utilises metaSPAdes assembler (Nurk et al. [2017]) to extract contigs from each sample independently. It can be useful in the case where assembly is planned to be performed during a study, as the assembly results are saved and can be reused. Next, the resulting contigs are transformed into features either directly or by combining them via de Bruijn graph construction.

Transition from graph components to the numerical features is done via k-mers alignment back to the components and calculation of depth and breadth coverage for each sample independently. The result of the feature extraction step is identical for all samples set of graph components used as features. For each sample the following steps are performed:

1. Take one component and split it into k-mers.
2. Make an intersection of component’s k-mers set and sample’s k-mers set.
3. Count the size of intersection and divide it by the size of the component. The resulting value from 0 to 1 is the *breadth* coverage.
4. The *depth* coverage is calculated by multiplying each common k-mer by it’s abundance in the sample. The resulting sum is divided by the size of the component to get positive real number – mean depth coverage.
5. Repeat the steps 1-4 for all components.

As a result, each sample is represented as two numerical vectors of length equal to the number of extracted graph components. The choice of vector for further use in classification models left to the user (by default *breadth* coverage is used).

Finally, all vectors are joined into the table of shape `N features` *×* `M samples`. Since the additional metadata for samples is not exploited during features construction (and generally not available, otherwise supervised algorithms should be preferred), the resulting feature table can be used for samples clustering and patterns detection.

### Supervised methods for feature extraction

Should the input dataset have the associated metadata to split samples into several categories, supervised methods can be applied to extract features specific for each group. Overall scheme of features extraction pipelines is presented in Algorithm 1 and on Supplementary Figure 2. MetaFX contains three independent strategies aimed at selection of specific k-mers for each group of samples, implemented in modules `unique`, `chisq`, and `stats`.

**Fig. 2.**
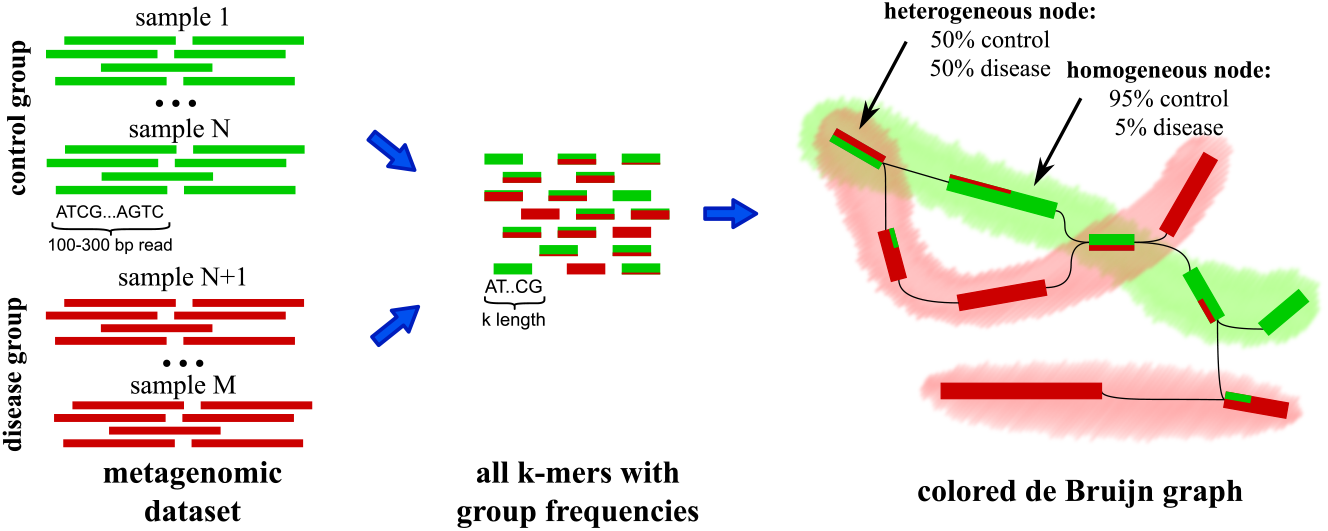
Workflow of `colored` feature extraction module. The extracted features are highlighted on de Bruijn graph with shades (green for control and red for disease group).

`Unique` algorithm searches for group-specific k-mers, defined as a k-mer present in at least *G* samples of certain group and absent in all other samples. *G* parameter can be chosen by researcher or automatically selected to obtain usable number of features (approximately 1000 to 10000) for each category.

`Chisq` and `stats` algorithms apply statistical tests to select group-significant k-mers. First method utilise chi-squared test for selection top significant k-mers, while second method combines chi-squared with Mann–Whitney *U* test to select k-mers with different occurrences between categories. Yates’s correction for continuity and Bonferroni correction for multiple comparisons are used.

All three aforementioned methods result in sets of k-mers specific for each category, that need to be transformed into features. Further, selected k-mers are used as pivots for local de Bruijn graph construction. Each category is processed independently. For a given category all samples are split into k-mers and are used to build de Bruijn graph. Next, selected specific k-mers are marked as starting points for search in the graph. For each starting point we perform the procedure of local search, based on combination of depth-first and breadth-first searches with early-stopping criteria. The number of branches studied during the search is defined by the launch parameter. The branches with no group-specific k-mers are discarded. As a result, numerous selected k-mers are grouped into graph components, reducing the resulting number of features. Calculation of numerical feature vectors for the components performed in the same manner as for the unsupervised methods. Finally, each component is saved into fasta file as non-overlapping contigs for further analysis and annotation.

#### Algorithm 1 Supervised feature extraction pipeline

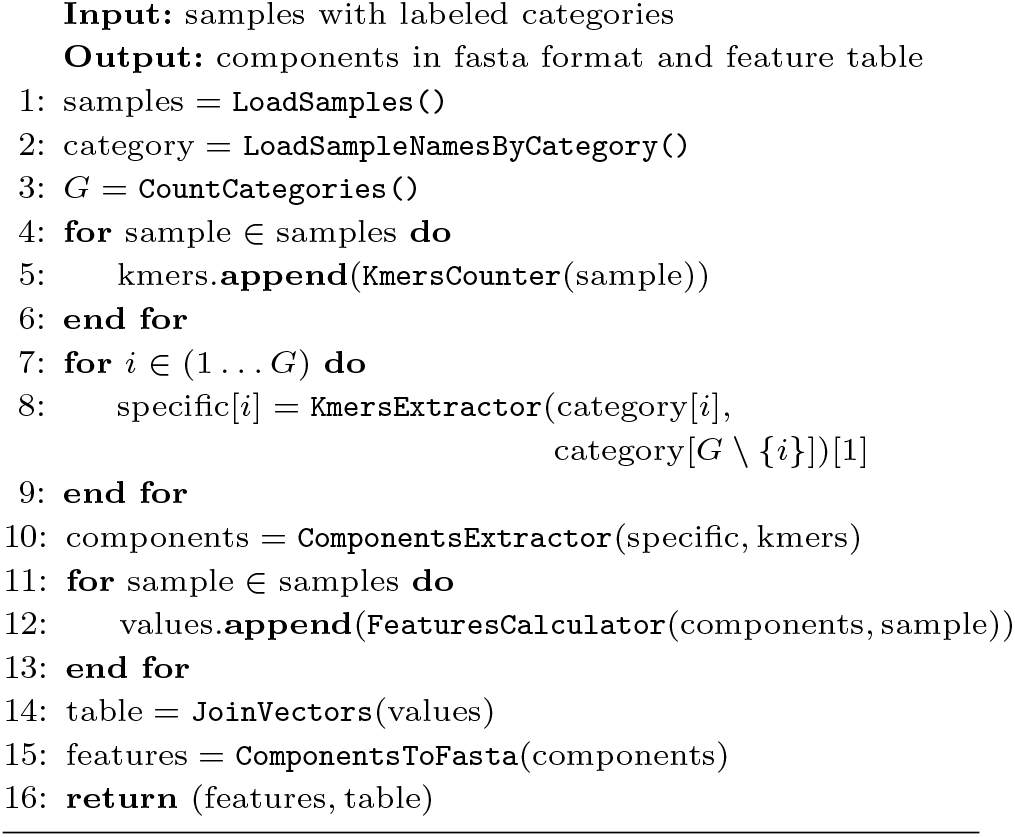

The last MetaFX module for supervised feature extraction is named `colored` and can process only two or three groups of samples (Figure 2). This limitation comes from the implementation details. However, the method is still useful for biomedical applications, e.g. comparing disease vs control group, or control vs treatment 1 vs treatment 2.

The distinctive advantage of the `colored` module is the control of the desired number of resulting features. Unlike the methods described above, it firstly assigns to each k-mer the probability vector to belong to each of the groups. The length of the vector equals to the number of categories. The value in vector is a ratio of number of samples of given category *G* containing the given k-mer to the total number of samples with that k-mer (see Equation 1).

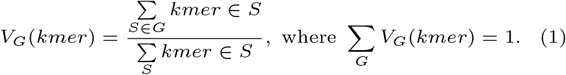

Further the global de Bruijn graph for all samples is constructed with nodes assigned `colors` based on k-mers probabilities. Finally, the colored graph is split into components. User can select to construct limited number of components for each group, starting from the group most probable k-mers, or to output all components around colored k-mers with probability above the threshold.

### Building predictive classification models

Having constructed the features, the next step is to analyse them. One option is to use the obtained feature table in machine learning algorithms. For that purpose several python scripts were developed. In case of the unsupervised feature extraction, MetaFX `pca` algorithm can be used to perform Principal Component Analysis (PCA) and visualisation of samples proximity. The implementation is based on scikit-learn library (Pedregosa et al. [2011]).

In case of the supervised feature extraction, `fit` or `cv` modules are used to train classification model and, optionally, tune the hyper-parameters. Tree-based models remain state-of-the-art classifiers for tabular data analysis (Grinsztajn et al. [2022]). Consequently, MetaFX supports Random Forest Classifier from scikit-learn library, XGBoost library for building tree-based gradient boosting models (Chen and Guestrin [2016]), and PyTorch library with custom neural network to showcase the possible analysis (Paszke et al. [2019]). However, the resulting feature table can be examined by any method of the researcher’s choice.

The constructed features and pre-trained classifiers are extremely valuable to speed up further research. Suppose the case of new study group of samples with unknown categories from the same type of the environment. Metafx `calc features` method should be called to count numeric feature values for new samples based on previously extracted graph features. This allows the researcher to skip the most resource-consuming steps of graph construction and feature extraction. Instead, basic k-mers manipulations enough to obtain numeric features for classification. After that, `predict` method can be used to classify novel samples with pre-trained model.

### Methods for data analysis and visualisation

Another option is to analyze features in-depth and visualise them. For convenient visual representation of extracted features, Bandage program (Wick et al. [2015]) was substantially improved and the required version is available at https://github.com/ctlab/BandageNG. MetaFX `bandage` module enables the joint visualisation of feature graphs and the trained Random Forest classifier (Supplementary Figure 3). Bandage web-interface is split into two screens: left for de Bruijn graph visualisation and right for Random Forest visualisation. Nodes in decision trees are annotated with the feature used as criterion. These features can be mapped back to the contigs of de Bruijn graph and highlighted. Thus we can detect the most used features in classification and analyse their graph environment.

**Fig. 3.**
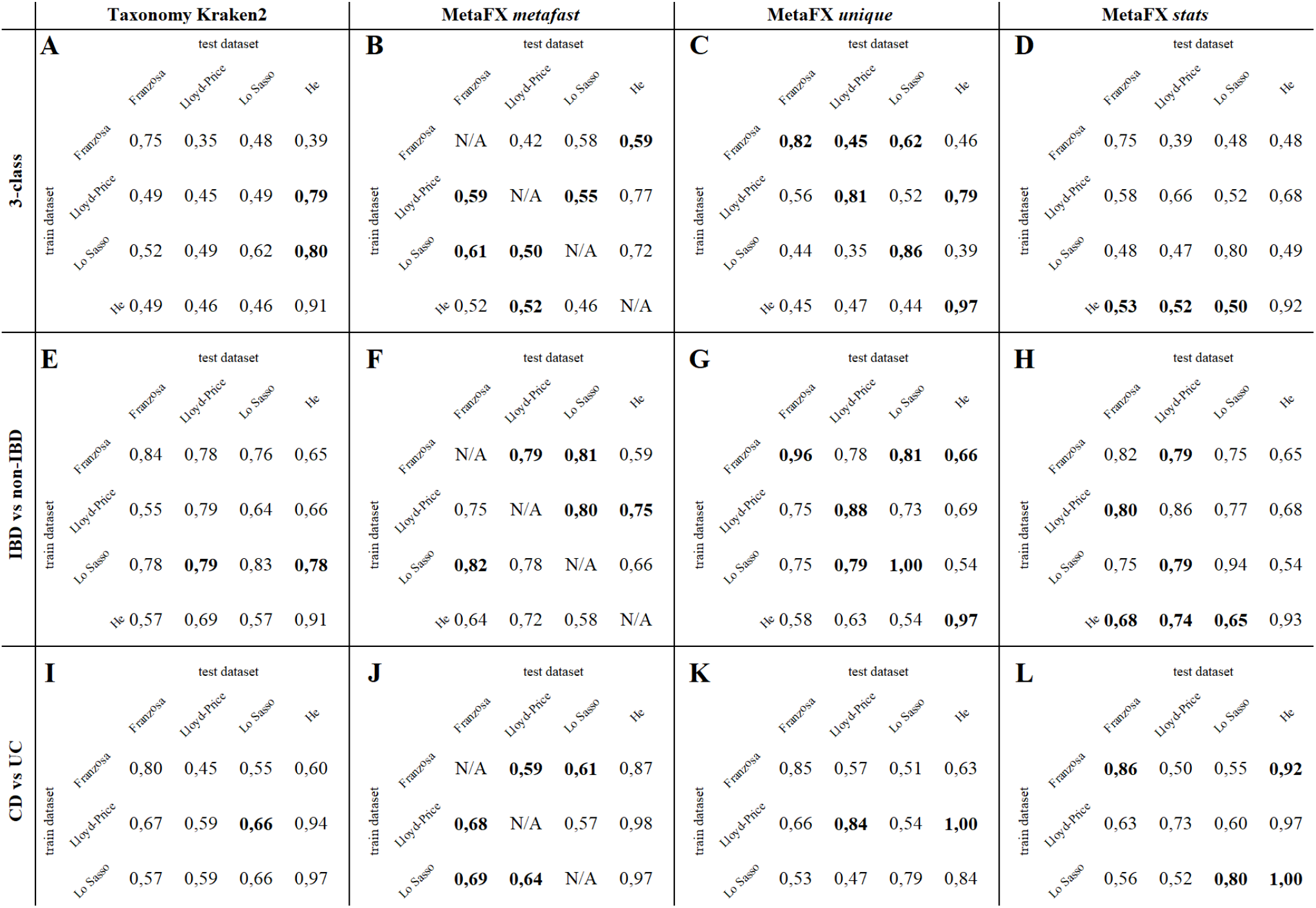
Inflammatory Bowel Disease Random Forest classification accuracy. Three scenarios were analysed: 3-class predictions (A-D), IBD vs non-IBD (E-H), CD vs UC (I-L). For each scenario for each train-test pair in bold highlighted one of four best methods. 5-fold cross-validation used for the same train-test pair, except for MetaFX `metafast` method (see details in text).

Metafx `feature analysis` module allows the detailed examination of the selected feature. Feature’s local environment is constructed for each sample via MetaCherchant toolkit (Olekhnovich et al. [2018]) and then all the environments can be simultaneously visualised in BandageNG. These visualisations can be manually explored for connections not obvious from contigs fasta files.

## Results

MetaFX was tested on real human gut metagenomic dataset of 590 samples (*∼*1.8 Tb of gzipped reads). Data was collected from 4 studies (Franzosa et al. [2019], Lloyd-Price et al. [2019], Lo Sasso et al. [2021], He et al. [2017]) focused on gut microbiome analysis of patients with inflammatory bowel disease (IBD). Samples are grouped into 3 categories: patients with Crohn’s disease (CD), patients with ulcerative colitis (UC) and control group of healthy individuals. Generalised information about datasets is present in Supplementary Table 1, per-sample statistics are available from authors’ original articles.

Four strategies were used for feature extraction. As a baseline, we performed taxonomic annotation of all samples via Kraken2 (Wood et al. [2019]) with *Standard* database of Refseq archaea, bacteria, viral, plasmid, human and UniVec Core collected on 14 March 2023 at Langmead 2023]. Annotations were filtered at species level and relative abundance was used as features in classification models.

Three other strategies are implemented in MetaFX. Firstly, reads were processed by splitting them into k-mers of length 31. Each sample was processed independently and all singletons were discarded. As a result, each sample is represented as a set of 31-mers. Next, we applied one unsupervised and two supervised methods for feature extraction.

MetaFX `metafast` unsupervised algorithm was applied to all samples in all datasets simultaneously, as we pretend the lack of corresponding diagnosis as metadata. MetaFX `unique` and `stats` methods were applied to each dataset to samples grouped by diagnosis. As a result of all three methods we obtained feature tables of relative breadth coverage of extracted features by each sample.

Further, we tackled three types of classification problems. Firstly, we aimed to predict one of three classes. Secondly, we aimed to distinguish between control samples and diseased samples. Finally, we aimed to differentiate between CD and UC diagnosis.

We applied Random Forest classification model with 100 decision trees and other default parameters from scikit-learn library (Pedregosa et al. [2011]). Each of four dataset was used as training, and the performance of classification model was estimated separately on other datasets. We also performed 5-fold cross validation of classifiers with the same train and test datasets. Since the feature sets for `metafast` method are the same in our experimental setup for all datasets, we did not perform the cross-validation. For CD vs UC problem, He at al. dataset was not used for training, as it lacks UC samples. The accuracy of classifications are shown on Figure 3.

For each type of classification problem (Figure 3 A-D, E-H, I-L) we selected the best feature extraction method for each pair of train-test datasets. In 3-class problem `unique` method showed the best in cross-validation cases, while `metafast` algorithm was the best in most other pairs. Taxonomic annotation outperformed other methods only once with Lo Sasso training dataset and He testing dataset. The increase in accuracy for the best method compared with taxonomic annotation ranges from 6 to 36 % for the same train-test pair, and from 1 to 20 % for different datasets. This fact can be explained by microbiome differences between cohorts characterised by lifestyle, dietary habits, etc. Additionally, we compared our results on Franzosa et al. [dataset with the classification accuracy from the original article. In cross-validation study, we reached 82 % accuracy by MetaFX `unique` module, while original article got only 65 % accuracy on taxonomic species.

In 2-class IBD vs non-IBD problem, the trend remains the same. For cross-validation `unique` algorithms shows the best results. However, `stats` feature extraction method perform better for different train-test pairs. Compared with 3-class problem, the increase in classification accuracy here is less pronounced due to a simpler task formulation. In 2-class CD vs UC problem, we also observe the increase in classification accuracy obtained with MetaFX features compared with taxonomic annotation.

Unfortunately, we cannot determine the best feature extraction method to serve all experimental setups and cohorts. Nevertheless, the proposed algorithms retrieve the information hidden in sequencing data and unavailable for classical reference-based tools. The combination of methods implemented in MetaFX toolkit can provide more accurate classification results and new insights based on unannotated features.

Finally, we searched the extracted components for relevant biological explanation of the obtained results. Component sequences, obtained with `metafx unique` method, were taxonomically annotated using Kraken2 (v. 2.1.3) with standard human and bacterial databases (Lu et al. [2022]). Visualisation of obtained results was performed using the pheatmap package for R (Kolde [2019]).

According to our analysis (Figure 4), there is a clear pattern of microorganisms composing the beneficial intestinal flora in a healthy state: *Faecalibacterium spp*., *Ruminococcus torques, Bifidobacterium adolescentis, Bifidobacterium longum, Coprococcus spp*. It is also characterised with direct producers of short-chain fatty acids (SCFAs), in particular butyrate – *Roseburia hominis, Anaerostipes hadrus, Faecalibacterium prausnitzii* (O’Riordan et al. [2022]), and bacteria implicated in processes linked to butyrate synthesis – *Ruthenibacterium lactatiformans, Blautia wexlerae, Dorea longicatena* (Becker et al. [2022]). In addition to producing enough mucus and preserving the epithelial barrier, butyrate is crucial for the metabolism of intestinal cells. These bacteria also have a potential anti-inflammatory effect induced by preventing the synthesis of several cytokines (Siddiqui and Cresci [2021]). In IBD a reduction in the number of butyrate-producing bacteria can dramatically intensify the inflammatory process and complicate the disease’s course.

**Fig. 4.**
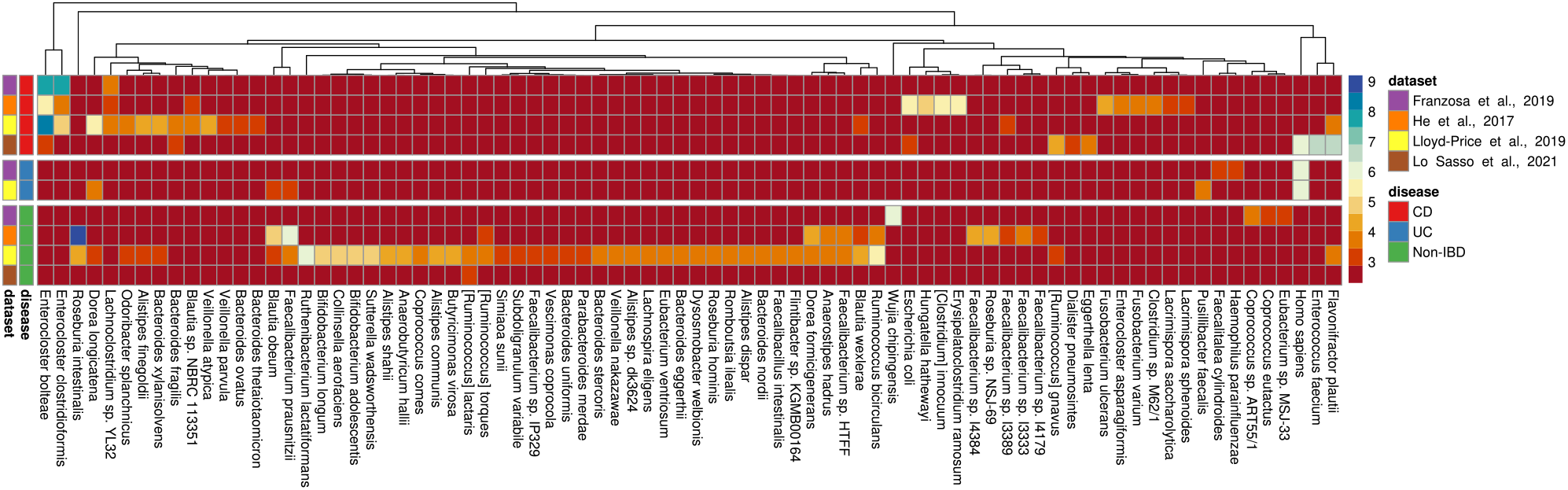
Taxonomic classification of components independently extracted via `metafx unique` algorithm and annotated for each dataset and category. Colour intensity calculated as logarithm of annotated components (blue shows high occurrence of taxa in dataset-category pair, red – low occurence).

Using MetaFX approach we detected several opportunistic pathogens associated with Crohn’s disease, including *E. coli, Hungatella hathewayi, Erysipelatoclostridium ramosum* and *Ruminococcus gnavus*. These pathogens are characteristic to the microbiota of the patients (Teh et al. [2021], Santiago et al. [2023], Ma et al. [2022]). It is also important to pay attention to *Clostridium innocuum*, which has been shown to translocate to the mesenteric adipose tissue and cause the processes of “creeping fat” formation and subsequent fibrosis. This can be a serious complication of the disease, because of the potential for creeping fat to invade the intestinal wall (Ha et al. [2020]). Furthermore, in each of the four analysed datasets, we found that Crohn’s disease was associated with a higher abundance of two *Enterocloster* bacteria: *Enterocloster clostridioformis* and *Enterocloster bolteae*. For instance, *E. bolteae* has the ability to produce microbially conjugated bile acids, which may have a negative effect on the disease (Guzior and Quinn [2021]).

In both Crohn’s disease and ulcerative colitis, there is a thinning of the mucous layer, massive inflammation and death of intestinal wall cells, which increases the release of DNA from dead cells. This explains the presence of human sequences in components even though they are absent from the healthy group.

Therefore, the developed MetaFX method helps to find nucleotide sequences associated with specific bacterial species highly correlated to the illness.

## Discussion

Metagenomic data is intensively examined for solving various problems of fundamental and applied science. However, existing methods analysing this type of data are imperfect, which can lead to the loss of critical information. Thus, there is a need to develop alternative techniques and approaches to improve the obtained results. At present, two paradigms of metagenomic data analysis can be distinguished: reference-based and reference free.

Reference-based methods include a wide range of applications for analysing metagenomes using reference sequences to perform taxonomic and functional annotation. Reference sequences may consist of gene sequences, metagenome-assembled genomes (MAGs), whole genomes, or parts of them. Overall, this type of analysis is limited by the completeness of databases and does not allow information on “microbial dark matter” to be included in the analysis. This may result in the loss of a crucial biological relation, which might be essential for machine learning models, among other things. Though, the addition of “lost” data may improve the efficacy of these models and contribute to a deeper understanding of the underlying biology of the issues in question.

Another group of methods for metagenomic analysis is reference-free methods. Firstly, these applications included genome-resolved metagenomic techniques. Discovering uncultivated microbial or viral diversity may be facilitated by recovering diversity directly from metagenomic data utilising genomes assembled from metagenomes. However, such pipeline is not always relevant when the researcher is faced with the task of quickly extracting information from metagenomes. Moreover, some of the useful information is also lost in the process of metagenome assembly and filtering out low-quality bins. Thus, there is a need to create algorithms that can process large datasets quickly but still work directly with the data without loss of information.

K-mer based approaches are alternative ways to analyse metagenomic data. They do not provide an in-depth understanding of microbial community structure, but allow for rapid exploratory data analysis to generate initial hypotheses. An additional advantage of this method is the inclusion of the unannotated part of the metagenome containing “microbial or viral dark matter”. These methods include MetaFast (Ulyantsev et al. [2016]) for rapid metagenome comparison, MetaCherchant (Olekhnovich et al. [2018]) for extracting the metagenomic environment of the target gene of interest, and RECAST (Olekhnovich et al. [2021]) for comparing donor and patient metagenomic sets in faecal transplantation experiments. The k-mer-based feature selection methods include approaches such as Commet (Maillet et al. [2014]), Simka (Benoit et al. [2016]), KMC3 (Kokot et al. [2017]), KmerGO (Wang et al. [2020]). Nevertheless, they do not allow further analysis of the identified features using taxonomic and functional annotation methods which makes biological interpretation of the results difficult.

To overcome these limitations, MetaFX, an open-source library for feature extraction from whole-genome metagenome sequencing data and classification of sample groups have been developed. MetaFX enables rapid extraction of features in the form of DNA sequences suitable for further study and generation of biological hypotheses. MetaFX can generate a limited number of features that are the best for samples classification, which is an added advantage of the proposed approach. Features can be extracted with or without samples metadata for supervised/unsupervised analysis. Additionally, MetaFX output can be used for classification models training.

The proposed approach has significant advantages because it eliminates the step of obtaining taxonomic or functional or other features to build models, and also preserves more useful information that can be used for prediction. Furthermore, tool aims to extract the most suitable information, including “dark matter”, which can be further annotated. It should be noted that MetaFX does not fully solve the problem of cross-cohort forecasting, alike many other classification methods. In other words, a classifier trained on one dataset does not always predict samples from another dataset well. The results are highly dependent on the similarity of samples between cohorts. The usefulness, benefits, and applicability of MetaFX were demonstrated in classification models for 590 metagenomes of patients with inflammatory bowel disease (IBD). IBD is still a serious and highly common group of diseases that lead to destructive changes in the digestive tract. All gastrointestinal tract processes, including inflammatory disorders, are directly influenced by the human gut microbiota. However, a thorough understanding of the precise mechanisms by which bacteria are involved in this process is still lacking. According to the obtained results, MetaFX produces superior disease prediction accuracy to the previous study by up to 17 % and improves classification results compared to taxonomic analysis by an average of 9 ± 10 %. As noted above, MetaFX generates features in the form of DNA sequences. They were assessed using taxonomic annotation methods and reproducible bacteria of the genus *Enterocloster* were found in samples from Crohn’s disease patients in all four studies. Bacteria of this genus can produce conjugated fatty acids which may have a negative effect on the course of the disease (Guzior and Quinn [2021]).

## Conclusion

Microbial communities often play crucial role in host environment and comparative metagenomics can shed light on the distinct microbiome properties. We have developed the MetaFX library to automate and simplify the process of extracting meaningful features from metagenomic datasets and training classification models for new samples analysis. MetaFX offers potential users a convenient and efficient reference-free workflow for analysing metagenomic sequencing data. MetaFX is capable of processing hundreds of samples within several hours and produces ready to annotation features as fasta contigs, that can be used both for bidimentional visualisation and for building predictive machine learning models. Moreover, the results can be explored visually by users thanks to the accompanying software BandageNG.

Applying additional taxonomic or functional annotation methods to MetaFX features results in generation of biological hypotheses based on data analysis of large numbers of metagenomic samples. Its possible future applications may include detection of disease-associated bacteria or functional parts of human gut microbiome in disease vs control studies, tracking the effect of certain medications or interventions on gut microbiota composition, and even screening and early diagnostics of possible illnesses based on pre-trained predictive models. Finally, MetaFX analysis is not limited to human gut and can be applied to other environments such as water, soil and cow rumen.

## Availability of source code and requirements

MetaFX source code is freely available on GitHub under the MIT license at https://github.com/ctlab/metafx. Detailed documentation is available at wiki page: https://github.com/ctlab/metafx/wiki. The tutorial for MetaFX based on mock microbial community analysis is available at https://github.com/ctlab/metafx/wiki/MetaFX-tutorial. Also a video with installation steps and basic instructions is available on YouTube: https://youtu.be/mTuP1jmOlI.

- Project name: MetaFX
- Project home page: https://github.com/ctlab/metafx
- Operating system(s): Linux/macOS
- Programming language: Java, Python
- Other requirements: JRE 1.8 or higher, Python 3.9.5 or higher, other requirements are available within the documentation
- License: MIT

## Data availability

All metagenomics data used in the current study were obtain freely from open databases. Details about samples and raw data can be found in the corresponding articles. Metagenomic sequences for Franzosa et al. [2019] are available from SRA BioProject PRJNA400072. Metagenomic sequences for Lloyd-Price et al. [2019] are available from SRA BioProject PRJNA398089. Metagenomic sequences for Lo Sasso et al. [2021] are available from the authors by the request. Metagenomic sequences for He et al. [2017] are available from the EBI database under the BioProject number PRJEB15371.

## Acknowledgment

We would like to thank Anastasiia Shostina for testing MetaFX and enthusiastic work on BandageNG tool and its integration with MetaFX. The authors thank the anonymous reviewers for their valuable suggestions.

## Author contributions statement

A.I. and V.U. developed the methodology and conception of the study; A.I. and V.P. designed, implemented and tested the software; E.O. validated the software; A.I. and E.O. collected the data for testing; A.I. and V.P. performed computational experiments; M.M. and E.O. performed biological analysis; A.I., M.M. and E.O. interpreted the results; E.O. and V.U. acquired the financial support; V.U. coordinated the project team; A.I. and E.O. wrote the manuscript; all authors reviewed and approved the final manuscript.

## Competing interests

The authors declare that they have no competing interests.

## Funding

Financial support for this study was provided by the Russian Science Foundation under the grant № 23-75-10125 https://rscf.ru/project/23-75-10125/. This work was performed using the core facilities of the Lopukhin FRCC PCM J. “Genomics, proteomics, metabolomics” (http://rcpcm.org/?p=2806).

## Notes

### Competing Interest Statement

The authors have declared no competing interest.

