## Supplementary figures and images for "MetaFX: feature extraction from whole-genome metagenomic sequencing data"

### Supplementary Figure 1

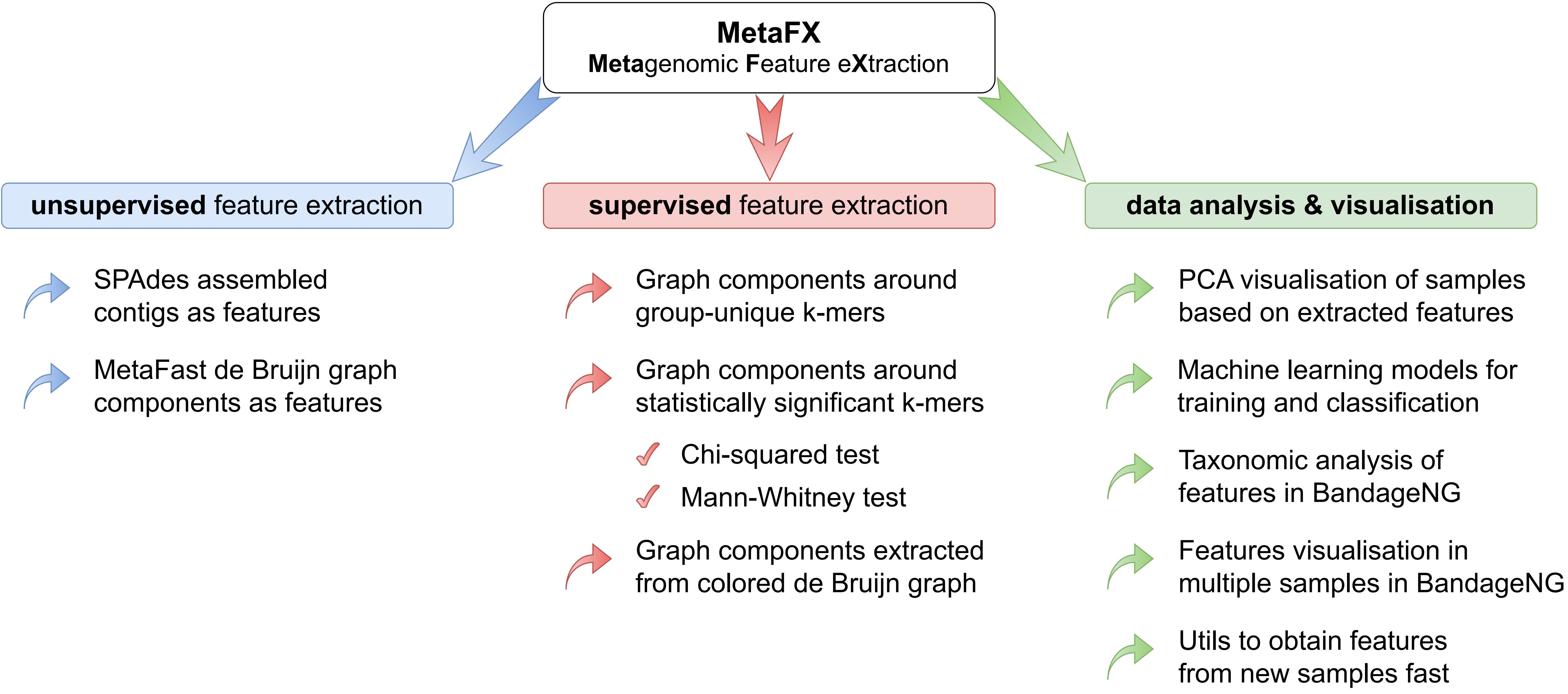

### Supplementary Figure 2

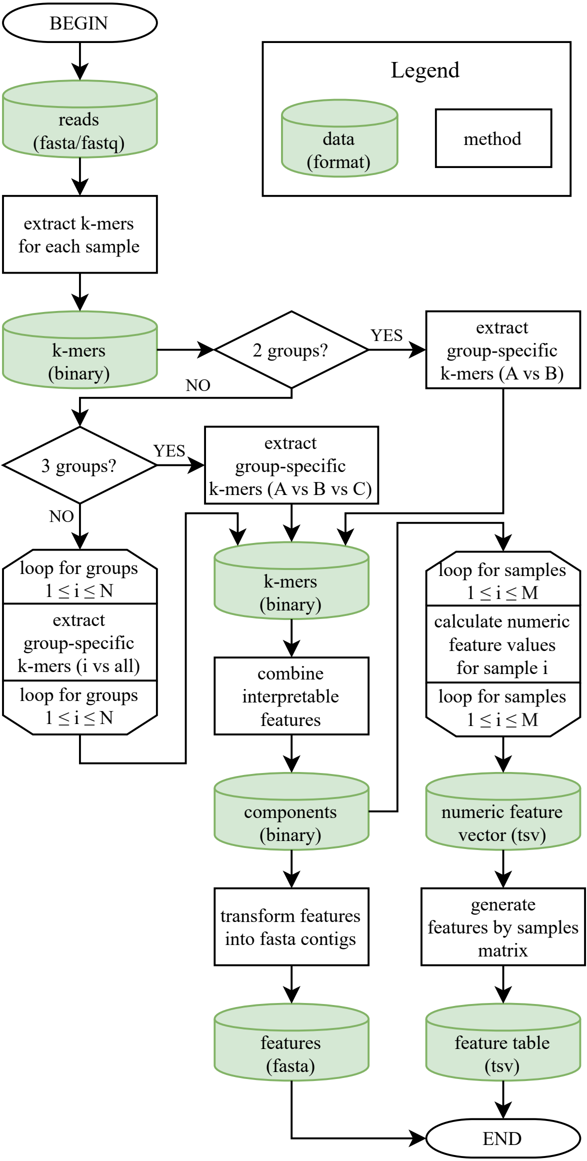

### Supplementary Figure 3

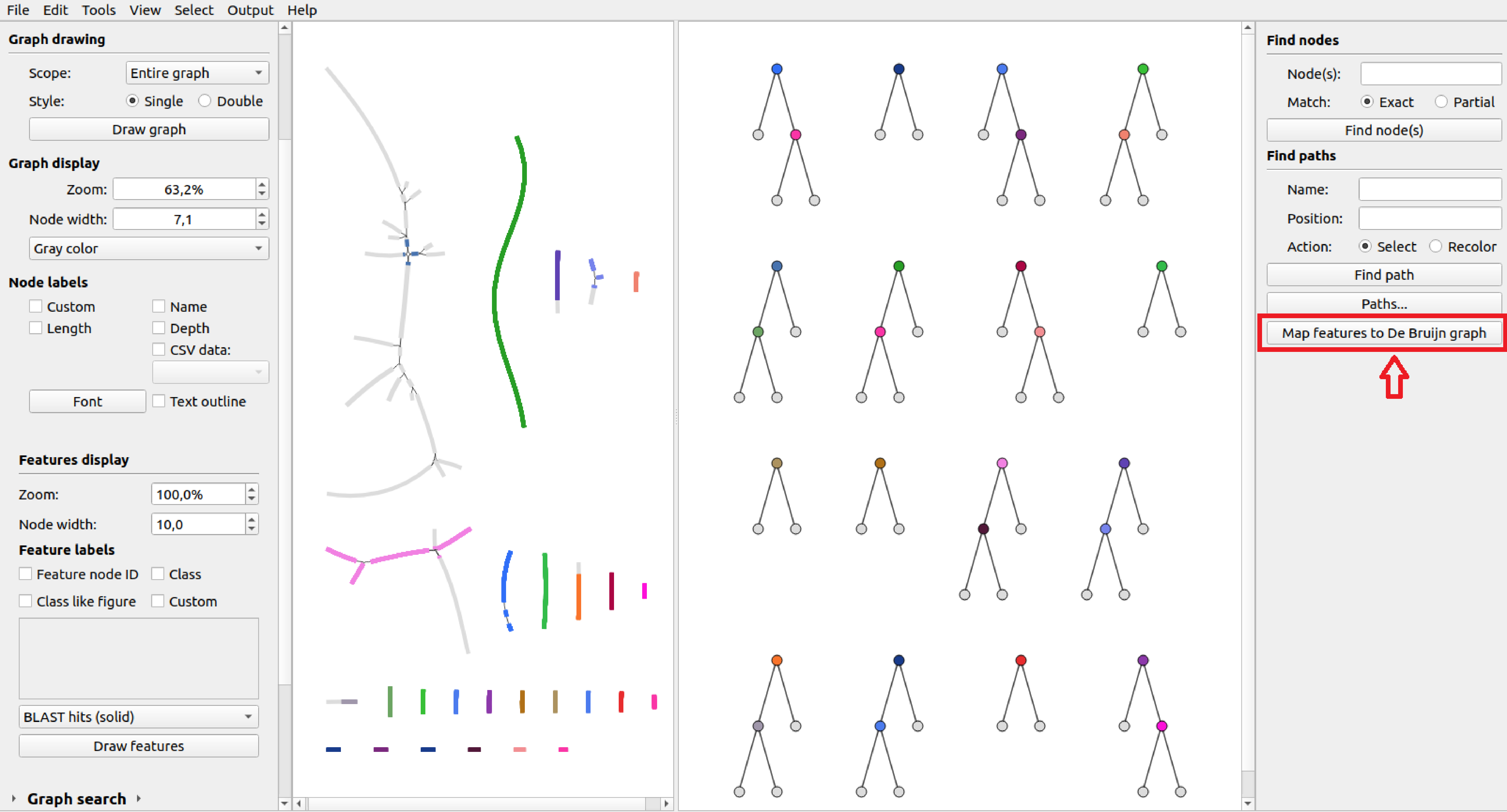
