## Supplementary Table 1 for "MetaFX: feature extraction from whole-genome metagenomic sequencing data"

**Table 1.** Datasets from human gut microbiome with inflammatory bowel diseases

| Dataset | Total samples | Crohn's disease | Ulcerative colitis | Control | # reads (mean±SD), mln | # k-mers (mean±SD), mln | # k-mers > 1 (mean±SD), mln |
|---|---|---|---|---|---|---|---|
| Franzosa et al. [2019] | 220 | 88 | 76 | 56 | 42.9 ± 33.3 | 490 ± 271 | 205 ± 105 |
| Lloyd-Price et al. [2019] | 130 | 65 | 38 | 27 | 24.3 ± 11.8 | 312 ± 152 | 129 ± 55 |
| Lo Sasso et al. [2021] | 124 | 40 | 42 | 42 | 55.3 ± 8.9 | 671 ± 343 | 256 ± 100 |
| He et al. [2017] | 116 | 63 | 0 | 53 | 55.8 ± 7.3 | 351 ± 123 | 179 ± 70 |
| **Total** | 590 | 256 | 156 | 178 | | | |
